# Molecular mechanism of regulation of the purine salvage enzyme XPRT by the alarmones pppGpp, ppGpp, and pGpp

**DOI:** 10.1101/2020.03.04.977603

**Authors:** Brent W. Anderson, Aili Hao, Kenneth A. Satyshur, James L. Keck, Jue D. Wang

**Author notes:** Correspondence, 1550 Linden Dr., 6478 Microbial Sciences Building, Madison, WI 53706. These authors contributed equally (Brent W. Anderson, Aili Hao).

## Abstract

The alarmones pppGpp and ppGpp mediate starvation response and maintain purine homeostasis to protect bacterial species. Xanthine phosphoribosyltransferase (XPRT) is a purine salvage enzyme that produces the nucleotide XMP from PRPP and xanthine. Combining structural, biochemical and genetic analyses, we show that pppGpp and ppGpp, as well as a third putative alarmone pGpp, all directly interact with XPRT and inhibit XPRT activity by competing with its substrate PRPP. Structural analysis reveals that ppGpp binds the PRPP binding motif within the XPRT active site. This motif is present in another (p)ppGpp target, the purine salvage enzyme HPRT, suggesting evolutionary conservation in different enzymes. However, XPRT oligomeric interaction is distinct from HPRT in that XPRT forms a symmetric dimer with two (p)ppGpp binding sites at the dimer interface. This results in two distinct regulatory features. First, XPRT cooperatively binds (p)ppGpp with a Hill coefficient of 2. Also, XPRT displays differential regulation by the alarmones as it is potently inhibited by both ppGpp and pGpp, but only modestly by pppGpp. Lastly, we demonstrate that the alarmones are necessary for protecting GTP homeostasis against excess environmental xanthine in *Bacillus subtilis*, suggesting that regulation of XPRT is key for regulating the purine salvage pathway.

## INTRODUCTION

The purine nucleotide GTP is a central molecule in DNA replication, RNA transcription, and protein translation. GTP can be synthesized either through an energetically costly *de novo* pathway or through efficient salvaging of preformed nucleobases. Substrate nucleobases are found in environments such as in exudates secreted by plant roots where they can be scavenged by soil bacteria for efficient nucleotide synthesis ^1,2^. The energetic benefit of the salvage reaction makes it the preferred pathway for GTP synthesis whenever nucleobases are available ^3,4^. However, excess intracellular GTP can lead to deleterious effects ^5^, and how the salvage pathway is regulated to protect organisms against external nucleobases fluctuations remains incompletely understood.

In bacteria, purine salvage can be regulated through inhibition of a key salvage enzyme, HPRT, by the nucleotide alarmones ppGpp and pppGpp ((p)ppGpp) ^5^. HPRT uses the essential metabolite phosphoribosyl pyrophosphate (PRPP) as a phosphoribosyl donor to convert the nucleobases guanine and hypoxanthine to GMP and IMP, respectively, which can then be converted to GTP (Figure 1A). HPRT is inhibited by (p)ppGpp competing with PRPP for binding the enzyme active site, which maintains GTP homeostasis under high guanine influx conditions ^6^.

**Figure 1.**
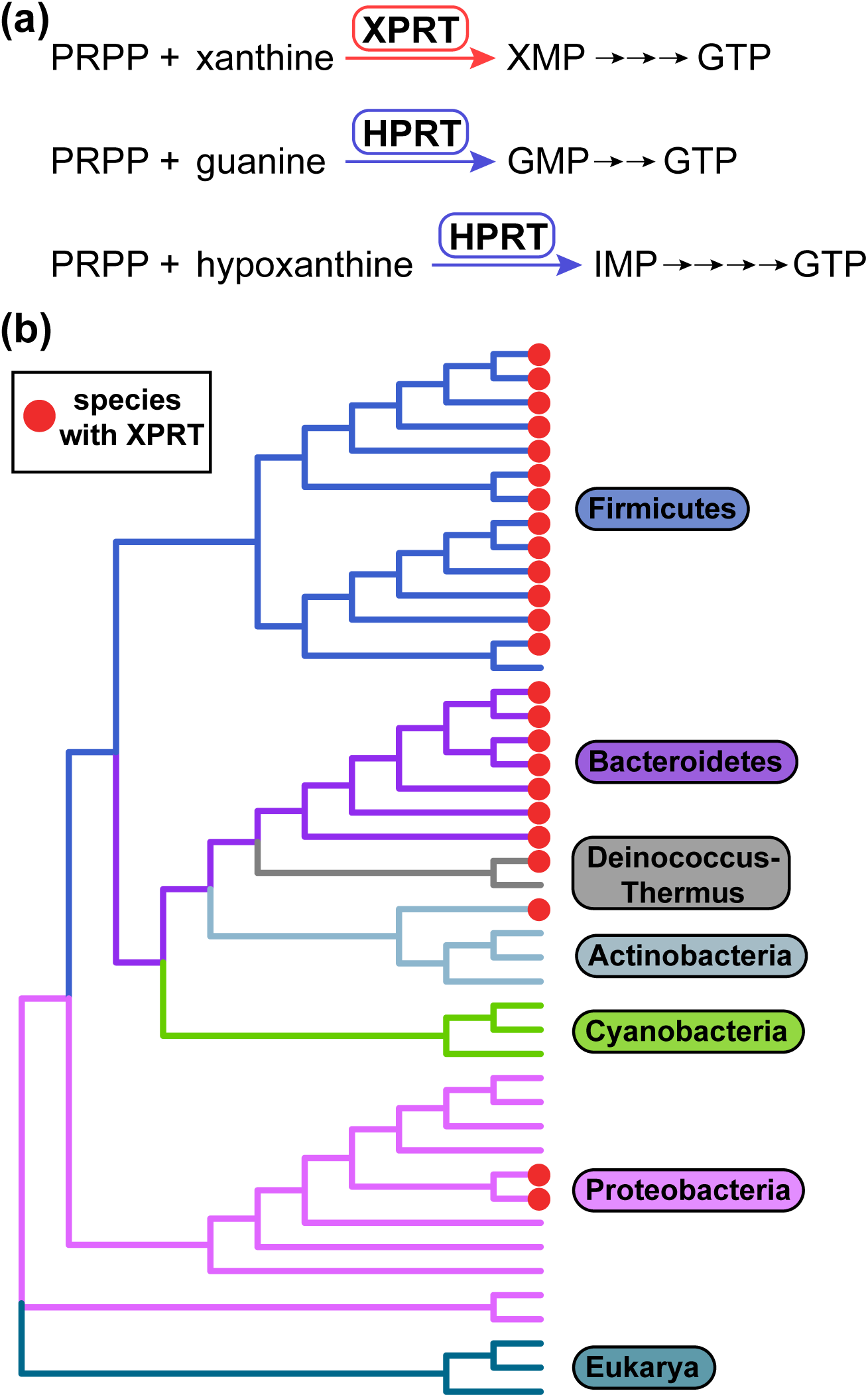
XPRT is a purine salvage enzyme conserved in Firmicutes and Bacteroidetes. **(a)** The purine salvage enzyme XPRT catalyzes the conversion of PRPP and nucleobase xanthine to the GTP precursor XMP. XPRT is homologous to the salvage enzyme HPRT that converts guanine or hypoxanthine to GMP or IMP. These salvage reactions are efficient pathways for GTP synthesis. (**b)** Cladogram constructed from 16S rRNA sequences from 41 bacterial species representing six bacterial phyla. Three eukaryotic 18S rRNA sequences included as the outgroup. Bacterial species containing XPRT are marked with red circles.

In the Gram-positive bacterium *Bacillus subtilis*, another phosphoribosyltransferase, XPRT, is also important for GTP synthesis. XPRT converts xanthine to XMP using PRPP as the phosphoribosyl donor (Figure 1A) ^7^. XPRT is part of the PRT protein family that use nucleobase substrates but has diverged, as demonstrated by a low sequence homology (10% identical) with HPRT. XPRT is conserved in the bacterial phyla Firmicutes and Bacteroidetes and is found in the phyla Actinobacteria, Deinococcus-Thermus, and Proteobacteria (Figure 1B). The lack of a mammalian XPRT counterpart also makes it a potential therapeutic target. However, how XPRT is regulated at an enzymatic level is poorly understood.

Here we reveal that XPRT is a regulatory target of (p)ppGpp as well as a target of the less well understood alarmone pGpp. Hereafter, we will refer to all three alarmones as (p)ppGpp unless otherwise noted. Similarly to HPRT, (p)ppGpp binds the XPRT active site using its PRPP binding motif to compete with substrate binding. However, the XPRT oligomeric interactions drastically differ from HPRT, resulting in unique regulatory features. First, (p)ppGpp binds an inter-subunit site comprised of residues from two monomers within an XPRT dimer resulting in cooperative binding of the regulatory ligand. Second, the XPRT binding pocket discriminates between pppGpp and ppGpp/pGpp through a flexible loop that covers the 5′ phosphate binding pocket, explaining why ppGpp and pGpp more strongly inhibit XPRT activity than pppGpp. Finally, (p)ppGpp appears necessary for GTP homeostasis maintenance upon exposure to excess external xanthine in *B. subtilis*.

## RESULTS

### pppGpp, ppGpp, and pGpp bind and inhibit the activity of XPRT

We first examined whether *B. subtilis* XPRT is a binding target of (p)ppGpp. To do so, we used the differential radial capillary action of ligand assay (DRaCALA) ^8^ to quantify the interaction between XPRT and ^32^P-labeled pppGpp and ppGpp. DRaCALA relies on the different migration properties of protein and ligand on nitrocellulose. Protein diffuses slowly but ligand diffuses rapidly when a solution is spotted on nitrocellulose. However, radiolabeled ligand interacting with protein co-migrates with the protein (e.g., Figure 2A). This interaction can be quantified as the fraction of total ligand bound with the protein. Using DRaCALA to obtain binding curves between XPRT and ^32^P-pppGpp and ^32^P-ppGpp, we found that *B. subtilis* XPRT is a binding target of pppGpp (K_d_ = 9.6 μM) and ppGpp (K_d_ = 0.95 μM) (Figure 2B). A third alarmone, pGpp, has been shown to be produced *in vitro* by a Firmicutes (p)ppGpp synthetase ^9^. We found that XPRT also binds pGpp (K_d_ = 0.76 μM) (Figure 2B). All three ligands interacted cooperatively with XPRT with Hill coefficients ∼ 2 for pGpp and ppGpp and ∼ 1.3 for pppGpp (Table 1).

**Table 1.**
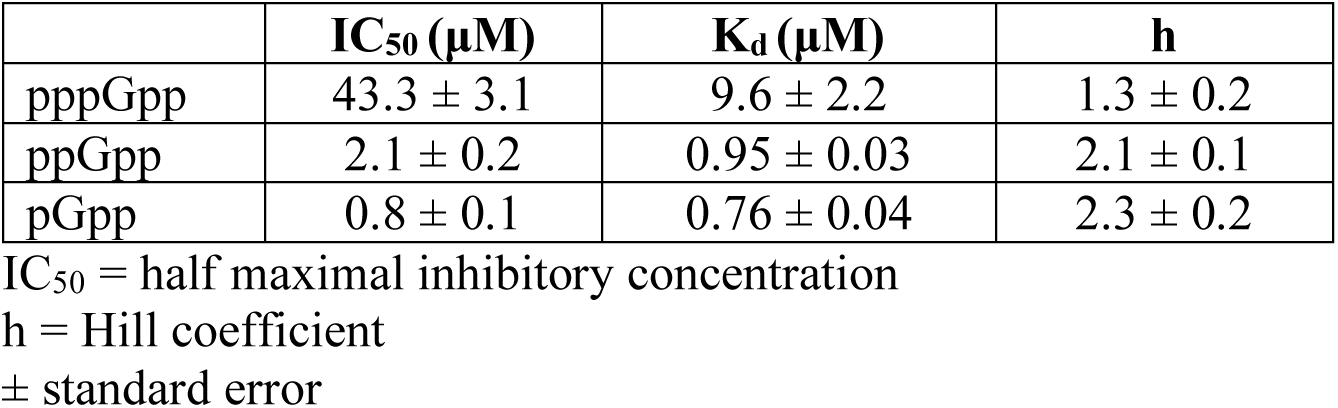
Parameters of alarmone interaction with XPRT

**Figure 2.**
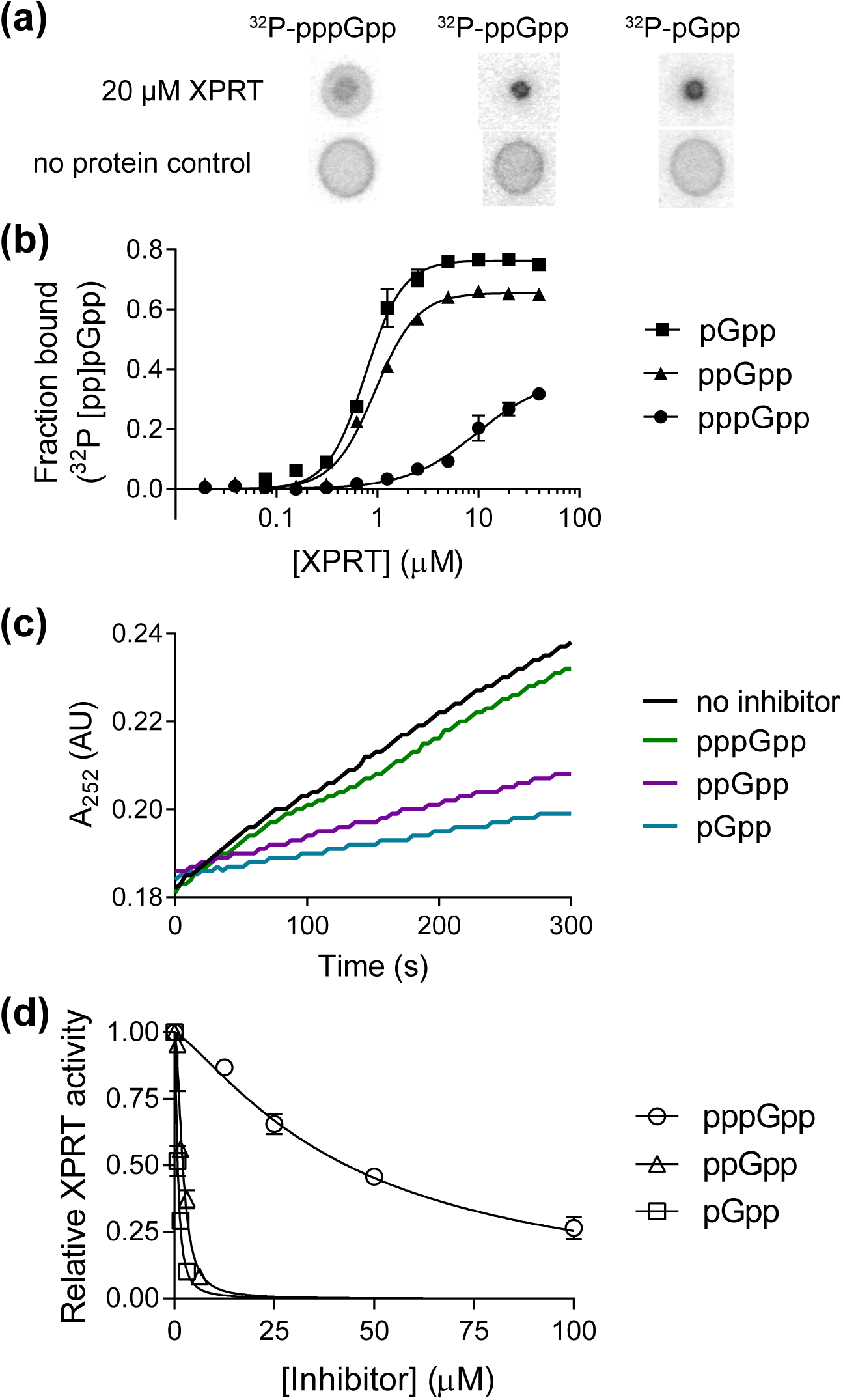
pppGpp, ppGpp, and pGpp bind and inhibit the activity of XPRT. **(a)** Representative DRaCALA data showing the interaction between 20 µM *B. subtilis* XPRT and ^32^P-pppGpp, ^32^P-ppGpp, and ^32^P-pGpp. The amount of radioactivity in the inner spot is a measure of the interaction. **(b)** Binding curves for ^32^P-pGpp, ^32^P-ppGpp, and ^32^P-pppGpp with varying XPRT concentrations. Binding determined by DRaCALA. Data are fitted to a single-site binding equation with the Hill coefficient. **(c)** Representative data showing first-order kinetic curves of XPRT activity with no inhibitor or with 1.56 µM pppGpp, ppGpp, or pGpp. Production of XMP measured as increased A_252_. Activity determined with 1 mM PRPP and 50 µM xanthine as substrates. **(d)** Inhibition curves showing that XPRT is more potently inhibited by pGpp and ppGpp than by pppGpp. See Table 1 for binding and inhibition parameters. Error bars represent SEM of triplicate.

To understand how these alarmones regulate XPRT, we tested their effect on XPRT activity. To measure XPRT activity, we used UV absorption (A_252_) to detect the rate of XMP production from the substrates xanthine and PRPP (e.g., Figure 2C), and initial velocities were calculated from these A_252_ curves. We found that pppGpp, ppGpp, and pGpp all inhibited XPRT activity (Figure 2C and 2D). ppGpp and pGpp potently inhibited activity with IC_50_ values of 2.1 μM and 0.8 μM, respectively (Figure 2D and Table 1). pppGpp was a weaker inhibitor with an IC_50_ value of 43.3 μM. The weaker inhibition by pppGpp corresponds with its reduced affinity for XPRT compared to ppGpp/pGpp (Figure 2B). This suggests that while all three alarmones regulate XPRT, it is differentially regulated by pppGpp and ppGpp/pGpp. Given that in cells the concentration of the alarmones can reach mM upon stress, these data demonstrate that XPRT is a regulatory target of pppGpp, ppGpp, and pGpp and all three alarmones inhibit XPRT activity.

### (p)ppGpp binds the conserved XPRT active site, and a flexible loop covering the pocket differentiates between pppGpp and ppGpp/pGpp

Next, we structurally examined the molecular mechanism of the (p)ppGpp-XPRT interaction. A structure of *B. subtilis* XPRT bound to ppGpp had already been deposited in the Protein Data Bank (PDB ID **<u>1Y0B</u>**), although the 5′-phosphates of ppGpp were not modeled in the deposited structure. Using the unmodeled F_o_-F_c_ difference density, we showed that there is sufficient density for a 5′-diphosphate in the ligand’s binding pocket (Figure 3A). Further refinement with ppGpp strongly supported the presence of a ppGpp molecule in the pocket (Figure 3B). The final structure is at 1.8 Å resolution with R_work_/R_free_ values of 0.160/0.208 (Table 2). XPRT-ppGpp crystallized as a tetramer in the asymmetric unit, but PISA analysis (Protein interfaces, surfaces, and assemblies’ service at European Bioinformatics Institute) predicts the biological unit to be a dimer (Figure 3C) ^10^.

**Table 2.** XPRT-ppGpp structure re-refinement statistics.

| **Refinement** |  |
| --- | --- |
| Resolution range (highest resolution bin) (Å) | 34.23 - 1.8 (1.864 - 1.8) |
| R<sub>work</sub>/R<sub>free</sub><sup>a</sup> (%) | 15.98 / 20.83 |
| r.m.s.<sup>b</sup> deviations |  |
| Bonds (Å) | 0.0038 |
| Angles (Å) | 0.76 |
| Ramachandran statistics (%) |  |
| Favored | 96.21 |
| Allowed | 3.66 |
| Disallowed | 0.13 |
| Rotamer outliers (%) | 0.00 |
| No. atoms |  |
| Macromolecules | 11763 |
| Ligands | 188 |
| Solvent | 756 |
| B factor (Å<sup>2</sup>) |  |
| Macromolecules | 25.98 |
| Ligands | 25.31 |
| Solvent | 33.25 |

<sup>a</sup>R<sub>work</sub>/R<sub>free</sub> = Σ||F<sub>obs</sub>| - |F<sub>calc</sub>||/|F<sub>obs</sub>|, where the working and free R factors are calculated by using the working and free reflection sets, respectively. The free R reflections were held aside throughout refinement. <sup>b</sup>Root mean square

**Figure 3.**
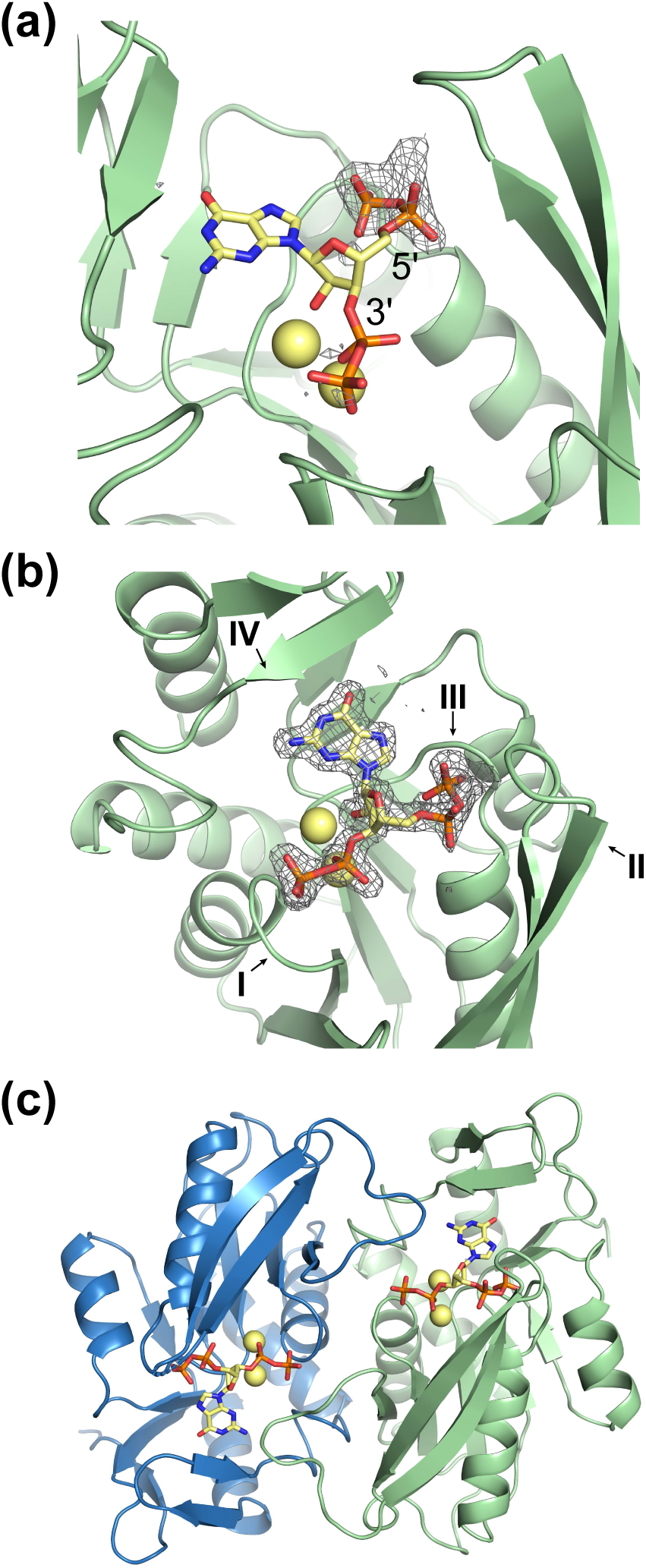
Costructure of *B. subtilis* XPRT and ppGpp. **(a)** Refinement of *B. subtilis* XPRT crystallized with ppGpp (PDB ID 1Y0B). The deposited structure lacked 5’-phosphates, but 5’-phosphates fit the F_o_-F_c_ difference density (contoured to 2.5 σ) at the 5’-carbon. In A-C, yellow spheres are Na^+^ ions crystallized with ppGpp. **(b)** ppGpp binding an XPRT monomer. Omit difference density shown contoured to 2.5 σ. I-IV refer to the four loops of the PRT active site. **(c)** Biological dimer of XPRT-ppGpp as predicted by PISA analysis. Each monomer in the dimer binds one ppGpp molecule.

The XPRT active site comprises four loops (I, II, III, IV) (Figure 3B). ppGpp binds this site in XPRT with its phosphates extending between loops I and III and the purine base positioned below loop IV (Figure 3B and 4A). Most residues interacting with ppGpp are contained within one monomer (Figure 4A). However, the second monomer in the XPRT dimer is also involved through a loop that extends over the ppGpp binding pocket (Figure 4A). We define this loop as the “bridging loop” since it bridges the monomer-monomer interface.

**Figure 4.**
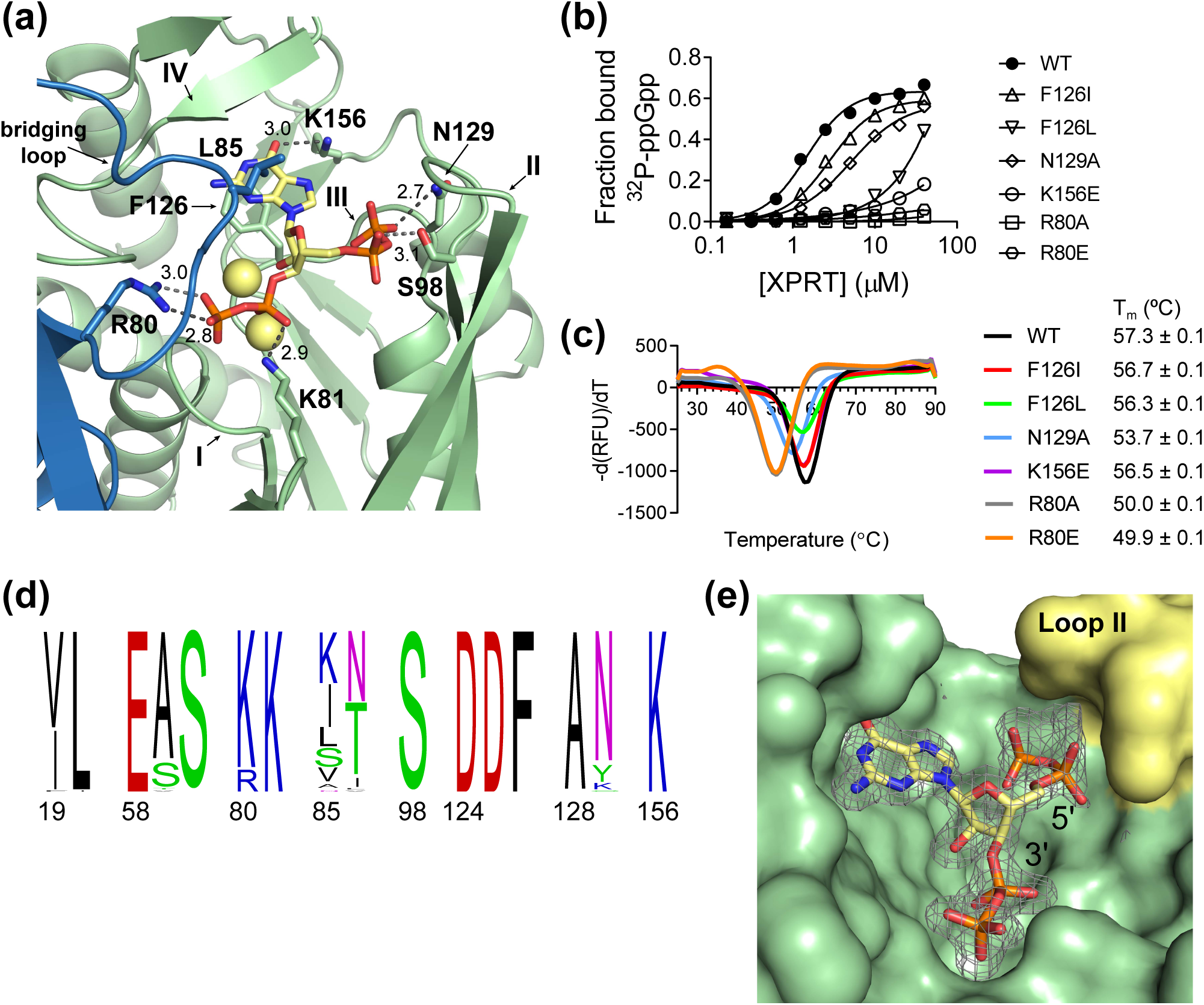
(p)ppGpp binds the conserved XPRT active site and a flexible loop covering the pocket differentiates between pppGpp and ppGpp/pGpp. **(a)** Select residues involved in the ppGpp-XPRT interaction. Hydrogen bonding shown as dotted lines and is measured in Å. The blue protein is the adjoining monomer in the structure. Yellow spheres are Na^+^ ions. **(b)** DRaCALA of *B. subtilis* XPRT active site variants shows that ^32^P-ppGpp binding is weakened by altering these residues. Points are mean of triplicate for all but F126L (duplicate). Error bars represent SEM for triplicate and range for duplicate. Error bars may be smaller than the height of the symbols. **(c)** Derivative curves from differential scanning fluorimetry of XPRT variants. Curves show the mean of triplicate reactions. Melting temperature (T_m_) is the mean of three replicates ± SEM. **(d)** Frequency logo of the ppGpp-binding residues from 62 bacterial XPRTs. Numbering is according to *B. subtilis* XPRT. Residues are colored according to their class. Logo generated using WebLogo (UC Berkeley). **(e)** Surface view of the ppGpp binding pocket on XPRT. Omit difference density for ppGpp is shown contoured at 2.5 σ. The compression around the 5’ phosphates caused by loop II (yellow) may be responsible for weaker interaction with pppGpp than ppGpp/pGpp.

The residues that interact with ppGpp include peptide backbone interactions of loops I and III that coordinate the 3′- and 5′-phosphates (Figure 4A and S1). XPRT Phe126 also forms π-stacking interactions below the guanine ring of ppGpp, Leu85 from the bridging loop forms a hydrophobic hood above the guanine ring, and Lys156 coordinates the exocyclic oxygen from the base (Figure 4A). XPRT Arg80, Lys81, Ser98, and Asn129 side chains all form hydrogen bonds with ppGpp’s phosphates (Figure 4A and S1).

To test whether these residues are indeed involved in their interaction with ppGpp *in vitro*, we created variants of Phe126, Asn129, and Lys156, and found that each variation resulted in a stably folded protein yet with weakened interactions with ^32^P-ppGpp (Figure 4B and 4C). We also created variants at Arg80, which interacts with ppGpp from the bridging loop (Figure 4A). These variants interacted very weakly with ^32^P-ppGpp, although the variants’ protein stability was also compromised (Figure 4B and 4C).

Since XPRT orthologs are conserved across Firmicutes and Bacteroidetes and are found in other bacterial phyla (Figure 1B), we examined the conservation of the binding pocket across these phyla. We aligned 62 bacterial XPRTs and constructed a frequency logo of the (p)ppGpp-interacting residues (Figure 4D). The binding pocket is strikingly conserved, with nine of the 16 residues being invariant (Glu58, Ser60, Lys81, Ser98, Asp124, Asp125, Phe126, Ala128, and Lys156). We conclude that the (p)ppGpp binding pocket on XPRT is conserved across bacterial XPRTs.

The XPRT-ppGpp structure provides an explanation for our earlier observation that ppGpp or pGpp are stronger inhibitors of XPRT than pppGpp (Figure 2C and D). In the structure, loop II forms a hood over the 5′-phosphate binding site (Figure 4E), compressing the pocket where the 5′-phosphates of pppGpp, ppGpp, and pGpp are positioned. Fewer 5′-phosphates in pGpp and ppGpp likely allow the inhibitor to better bind the active site and potently inhibit enzyme activity.

In summary, pppGpp, ppGpp, and pGpp bind the conserved XPRT active site, and XPRT displays differential regulation by pppGpp and ppGpp/pGpp due to a loop conformation that compresses the 5′-phosphate binding site.

### (p)ppGpp competes with PRPP to inhibit XPRT

We had previously obtained the co-structure of (p)ppGpp with the purine salvage enzyme HPRT ^6^. Overlaying the structures of ppGpp-XPRT and ppGpp-HPRT ^6^ shows that both ligands bind the enzyme active sites in a similar conformation (Figure 5A). We also overlaid the XPRT-ppGpp complex with HPRT bound to substrates PRPP and 9-deazaguanine (an inactive guanine analog) (Figure 5B) ^6^. Notably, ppGpp overlaps significantly with the substrates. The guanine ring of ppGpp likely binds in a position similarly to xanthine. The 5′-phosphates of ppGpp and PRPP share the same pocket, and the 3′-phosphates of ppGpp overlap with the 1′-phosphates of PRPP (Figure 5B).

**Figure 5.**
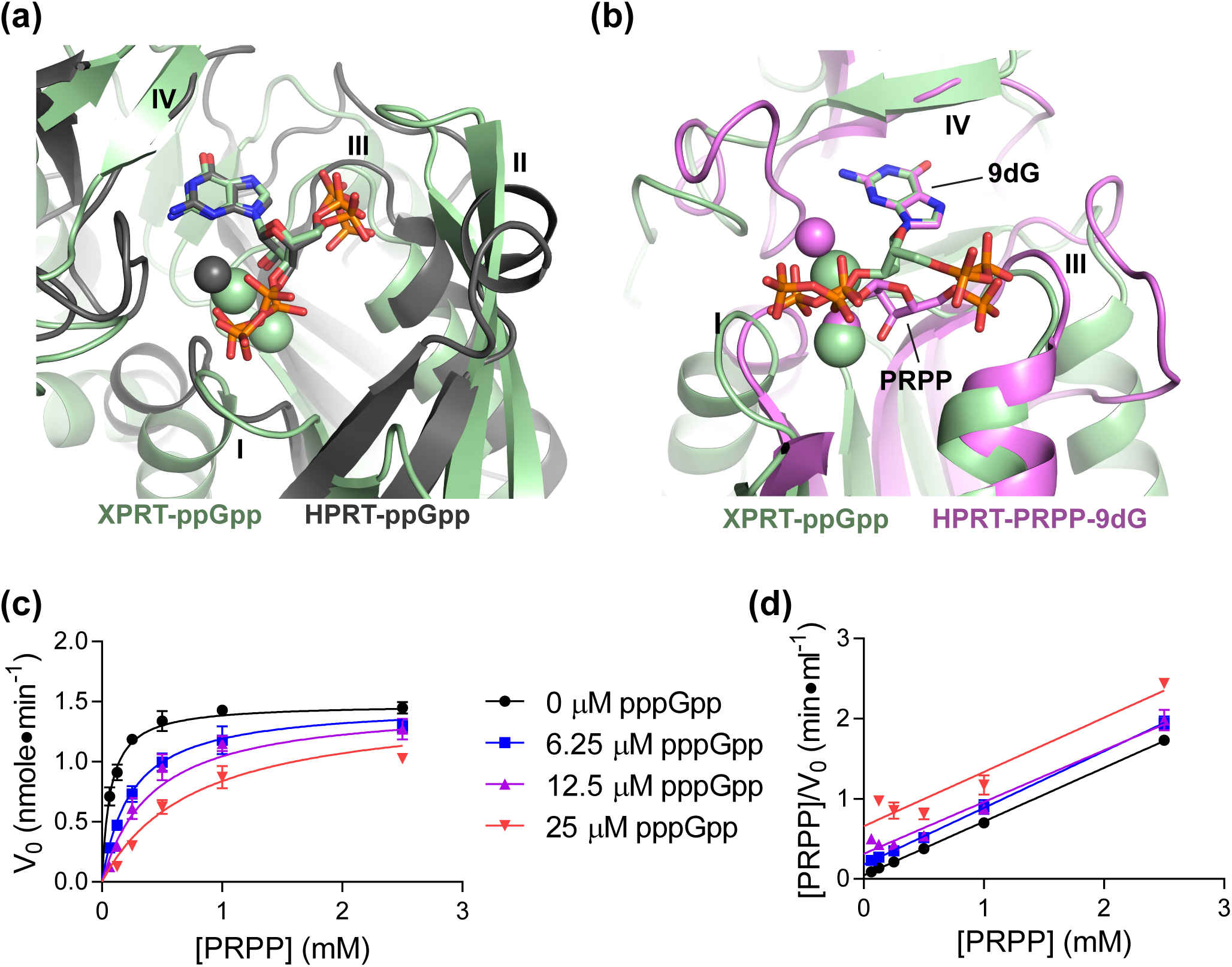
pppGpp competes with PRPP to inhibit XPRT. **(a)** Overlay of XPRT-ppGpp (green) and ppGpp-bound *B. anthracis* Hpt-1 (gray; PDB ID 6D9S). Alignment based on ppGpp molecules. Gray sphere is Mg^2+^ crystallized with HPRT-ppGpp. Green spheres are Na^+^ crystallized with XPRT-ppGpp. **(b)** Overlay of XPRT-ppGpp (green) and substrates-bound *B. anthracis* Hpt-1 (pink; PDB ID 6D9R). Alignment based on guanine of ppGpp and 9-deazaguanine (9dG) of substrates. ppGpp binds the XPRT active site and overlaps with substrate binding. Loop II for both proteins hidden for clarity. Pink spheres are Mg^2+^ crystallized with HPRT-ppGpp. Green spheres are Na^+^ crystallized with XPRT-ppGpp. **(c)** Initial velocities of *B. subtilis* XPRT at varied pppGpp and PRPP concentrations. Data are fitted to a global competitive inhibition model with a K_i_ of 2.5 µM. **(d)** Hanes-Woolf transformation from the data on the left. Equivalent slopes indicate equivalent maximum velocities at each pppGpp concentration. Error bars represent SEM of at least three replicates.

Next, we examined whether (p)ppGpp competes with PRPP to inhibit XPRT using steady-state kinetics. We measured initial velocities of XPRT enzymatic reaction by examining the rate of synthesis of XMP at varied pppGpp and PRPP concentrations (Figure 5C). The data best fit a global competitive inhibition model, demonstrating that (p)ppGpp competes with PRPP (Figure 5C and 5D). In addition, the kinetic data yielded a K_i_ for pppGpp of 2.5 μM and a K_m_ for PRPP of 69 μM. Together with the structure, these data validate XPRT as a (p)ppGpp target and show that (p)ppGpp competes with PRPP for binding the active site to inhibit the enzyme.

### (p)ppGpp stabilizes XPRT dimers by establishing electrostatic interactions bridging the monomer-monomer interface

XPRT is an unstable dimer with a tendency to dissociate to monomers at low concentrations ^11^. Although the majority of (p)ppGpp binding residues are in the conserved PRPP binding pocket within a single XPRT monomer, both monomers in an XPRT dimer are involved in binding each molecule of ppGpp (Figure 4A and 6A). The second monomer contributes the bridging loop (residues 80-89) to the ppGpp binding site (Figure 6B). Leu85 in this loop creates a hydrophobic hood for the guanine ring of ppGpp (Figure 4A). The bridging loop also contains Arg80 and Lys81 (Figure 6B). Lys81 contacts the 3′-phosphates of the ppGpp within its own monomer, while Arg80 reaches across the monomer-monomer interface to contact the 3′-phosphates of ppGpp in the adjoining monomer (Figure 6B). In this way, the presence of (p)ppGpp creates a network of electrostatic interactions bridging the XPRT monomer-monomer interface.

**Figure 6.**
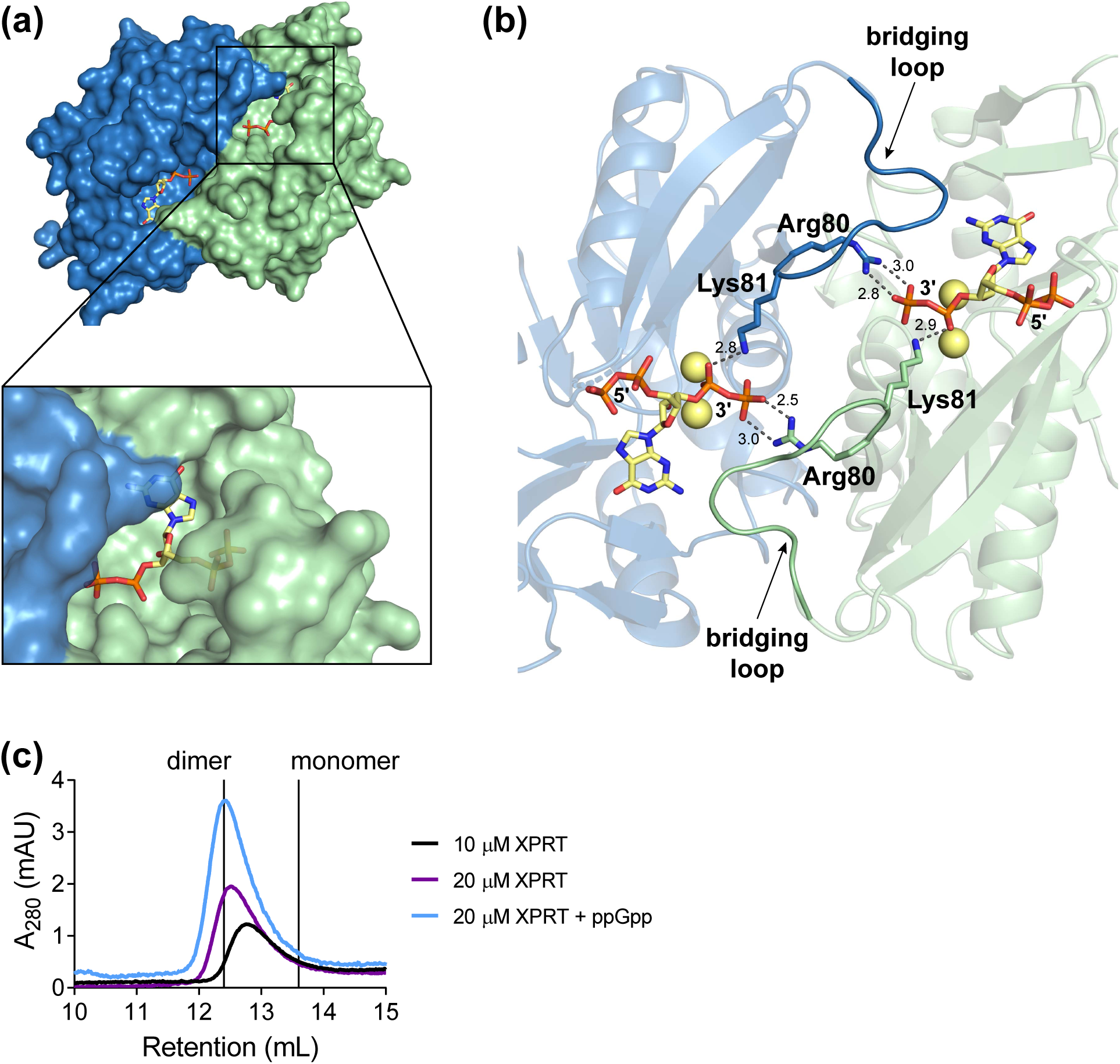
(p)ppGpp stabilizes XPRT dimers by establishing electrostatic interactions bridging the monomer-monomer interface. **(a)** *B. subtilis* XPRT dimer bound to ppGpp. Inset shows how ppGpp is covered by the second monomer. **(b)** Arg80 and Lys81 interact with the 3’-phosphates of ppGpp to create a network of electrostatic interactions across the monomer-monomer interface. Residues 80-89 comprise the bridging loop (opaque) that interacts with ppGpp across the interface. Spheres represent Na^+^ ions. Dashed lines represent hydrogen bonds measured in Å. **(c)** Size exclusion chromatograms of *B. subtilis* XPRT at 10 or 20 µM. ppGpp was added in the mobile phase. Vertical lines show the predicted retention volumes of monomeric and dimeric XPRTs based on their molecular weight (see Materials and Methods). The higher absorbance with ppGpp is likely due to ppGpp absorbing at A_280_.

Based on this structural evidence, we propose a model to explain the cooperative ligand binding we observed (Figure 2B): ppGpp binding to one site in an XPRT dimer would increase the affinity for ppGpp binding to the second site. This led us to predict that (p)ppGpp stabilizes the XPRT dimer by providing additional interactions for holding the monomers together. With size exclusion chromatography, we found that at a lower concentration (10 μM) of apo XPRT eluted at a molecular weight between monomer and dimer in agreement with previous reports (Figure 6C) ^11^. At a higher concentration (20 μM), XPRT eluted as a higher molecular weight although still not a stable dimer (Figure 6C). However, ppGpp in the mobile phase stabilized the dimeric XPRT population (Figure 6C). These data support that (p)ppGpp can stabilize the interaction between two XPRT monomers.

### (p)ppGpp protects against excess environmental xanthine

XPRT converts the purine base xanthine to the GTP precursor XMP. Xanthine is a nutrient that can be imported from environment by xanthine permease. In *B. subtilis*, XPRT is encoded by *xpt*, located in the same operon as xanthine permease, and is transcriptionally regulated by a purine-sensing riboswitch. XPRT has been shown to be expressed at copy numbers of ∼1,600-14,500 per cell across a variety of conditions, which allows the cell to effectively salvage xanthine (Figure 7A) ^7,12–14^.

**Figure 7.**
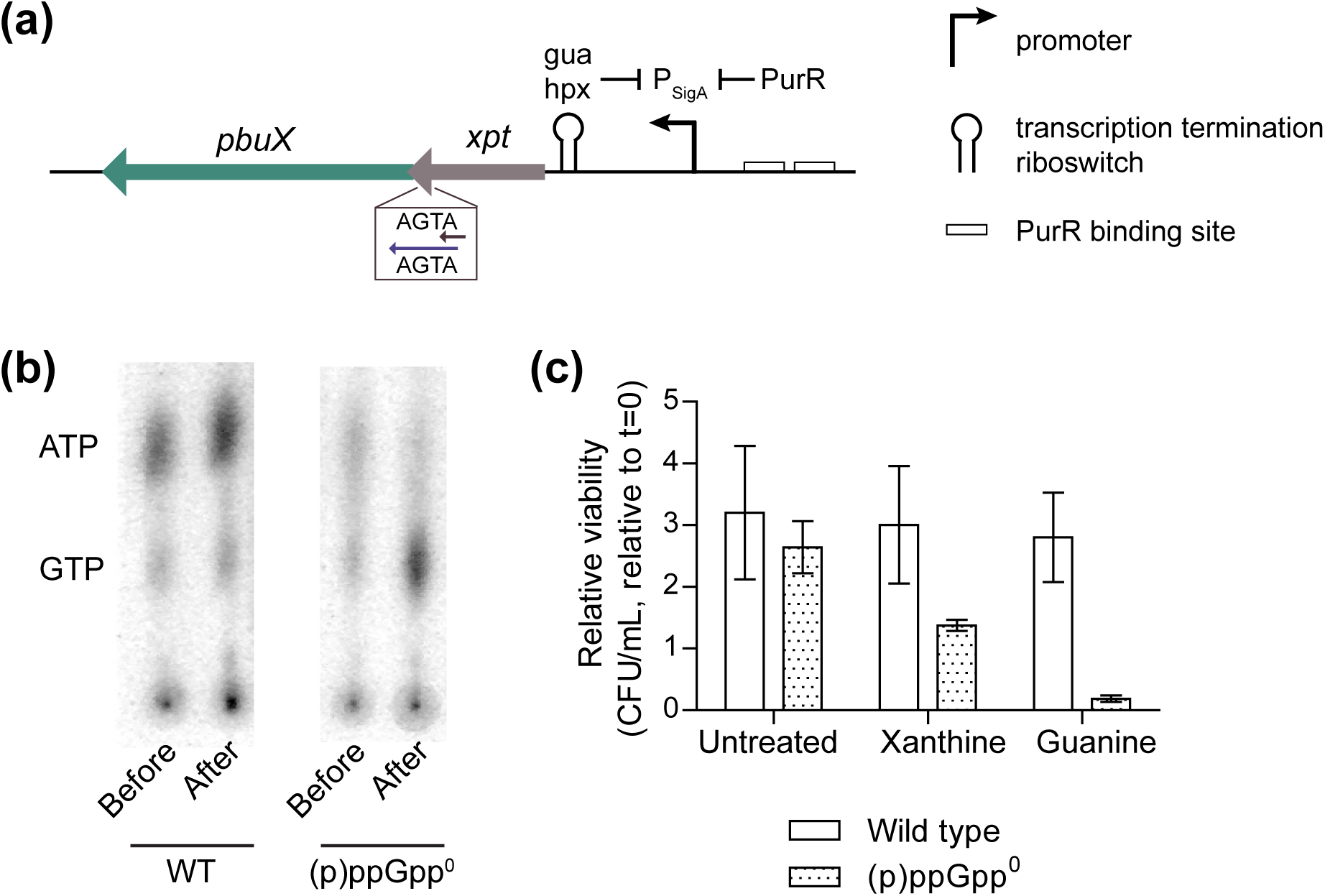
(p)ppGpp protects GTP homeostasis against excess environmental xanthine. **(a)** Genomic locus of *xpt*, which encodes XPRT. The *xpt* gene is in the same operon as *pbuX*, which encodes the xanthine permease. Gene expression is controlled by the purine repressor PurR and a transcription attenuation riboswitch that binds hypoxanthine (hpx) and guanine (gua). **(b)** Thin layer chromatography showing GTP and ATP in ^32^P-orthophosphate-labeled wild type and (p)p-pGpp^0^ *B. subtilis* before and after 30 minutes of treatment with 1 mM xanthine. Data are representative examples taken from the same TLC plate. **(c)** Viability of wild type and (p)ppGpp^0^ *B. subtilis* with 1 mM xanthine or 1 mM guanine. Sterile water was added to untreated sample. Cultures were grown in a minimal glucose medium with VILMTE (amino acids) to logarithmic phase prior to treatment for one hour, and colony forming units (CFUs) were calculated before and after treatment.

Since *B. subtilis* is capable of salvaging xanthine for GTP synthesis, we sought to understand the importance of (p)ppGpp in regulating xanthine utilization. We added xanthine to wild type and (p)ppGpp-null ((p)ppGpp^0^) *B. subtilis* growing exponentially in a defined glucose medium supplemented with the amino acids VILMTHRWE to support the growth of (p)ppGpp^0^. We then assessed GTP levels by thin layer chromatography. We found that GTP levels from xanthine-treated (p)ppGpp^0^ *B. subtilis* were elevated relative to wild type (Figure 7B). Higher GTP levels can promote growth, but too much GTP has been associated with reduced fitness of *B. subtilis* ^5^. We next examined the effect of xanthine on *B. subtilis* fitness by measuring colony forming units before and after xanthine addition. We found that in a (p)ppGpp^0^ *B. subtilis* strain, the addition of xanthine reduced growth compared to wild type cells (Figure 7C). As a control, we also added the nucleobase guanine to (p)ppGpp^0^, and it reduced viability as previously observed ^5^ (Figure 7C). These data reveal that excess environmental xanthine can disrupt GTP homeostasis and viability and (p)ppGpp is necessary to protect GTP homeostasis.

## DISCUSSION

(p)ppGpp is known to interact with multiple cellular targets, both at induced levels and basal levels, to protect cells against stress and maintain homeostasis. Here we combined structural and biochemical analysis to characterize a new (p)ppGpp target, the enzyme XPRT. We showed that (p)ppGpp binds to its active site at a PRPP binding motif and competes with substrate binding. We identified several unique features of regulation of XPRT due to its distinct oligomeric interactions, resulting in cooperativity and differential selectivity of alarmones. Our data indicate that basal levels pppGpp, ppGpp, and pGpp potently regulate XPRT activity to maintain GTP homeostasis. Combined with our previous results that (p)ppGpp inhibits the purine salvage enzyme HPRT, these data strengthen the model that basal (p)ppGpp protects GTP homeostasis against excess environmental purines by gating all purine entry points through inhibition of salvage enzymes.

### A common theme: PRPP motif in phosphoribosyltransferases binds (p)ppGpp

In recent years, many (p)ppGpp targets have been discovered, raising an important question of whether there are common themes in (p)ppGpp binding sites. While one common theme among many targets is that (p)ppGpp binds GTP binding sites (e.g., DNA primase and GTPases ^15,16^), we recently found that (p)ppGpp binds a PRPP binding motif, rather than at a GTP binding motif, in HPRT ^6^. In this work, we show that XPRT is another (p)ppGpp target that shares this PRPP binding motif, suggesting that the PRPP binding motif may be another common theme among many targets not harboring a GTP-binding site. This is not surprising given the structural similarity between the ribose and phosphates of PRPP and ppGpp, which would suggest that the PRPP binding motif is capable of interacting with (p)ppGpp.

XPRT is part of the PRT protein family that use nucleobase substrates but has diverged, as demonstrated by its low sequence homology with either *E. coli* XGPRT (12% identical) or *B. subtilis* HPRT (10% identical). The *B. subtilis* XPRT paralog specifically uses xanthine as a substrate, while the *E. coli* XGPRT paralog uses both xanthine and guanine as substrates ^11,17^. Orthologs to *B. subtilis* XPRT are found across Firmicutes and Bacteroidetes with additional representatives in Deinococcus-Thermus, Actinobacteria, and Proteobacteria (Figure 1B). The conservation of the (p)ppGpp binding site suggests that it is likely that XPRTs across these bacteria are also regulated by (p)ppGpp (Figure 4C). XPRT was not identified in a recent screen for (p)ppGpp binding proteins in the Firmicute *S. aureus*, but the screen was limited to ∼85% of the *S. aureus* proteome ^18^. Further work is needed to verify that (p)ppGpp interacts with XPRT from this and other organisms.

There are several other recently identified (p)ppGpp targets that contain the PRPP binding motif, including the phosphoribosyltransferases XGPRT, UPRT, and PurF from *E. coli* ^6,19,20^. All these enzymes bind PRPP as a substrate. However, not all phosphoribosyltransferases bind (p)ppGpp with the PRPP motif. The amidophosphoribosyltransferase PurF from *E. coli*, which converts PRPP and glutamine to phosphoribosylamine, binds ppGpp at an allosteric site away from the PRPP-binding active site ^20^. It will be interesting to examine whether XGPRT and UPRT from *E. coli* are inhibited through (p)ppGpp binding the PRPP site. If so, this will support a model that (p)ppGpp regulates phosphoribosyltransferases with nucleobase substrates.

### Variation: Oligomeric interaction affects cooperativity, affinity, and specificity

Despite HPRT and XPRT sharing a common PRPP- and (p)ppGpp-binding motif, they display striking differences in how they interact with (p)ppGpp due to distinct oligomeric interactions (Figure 8A). In contrast to the XPRT dimer that binds (p)ppGpp, HPRT binds (p)ppGpp as a tetramer (dimer-of-dimers) (Figure 8B and 8C), and the differences in (p)ppGpp binding arise from diversification of oligomeric interfaces in each protein.

**Figure 8.**
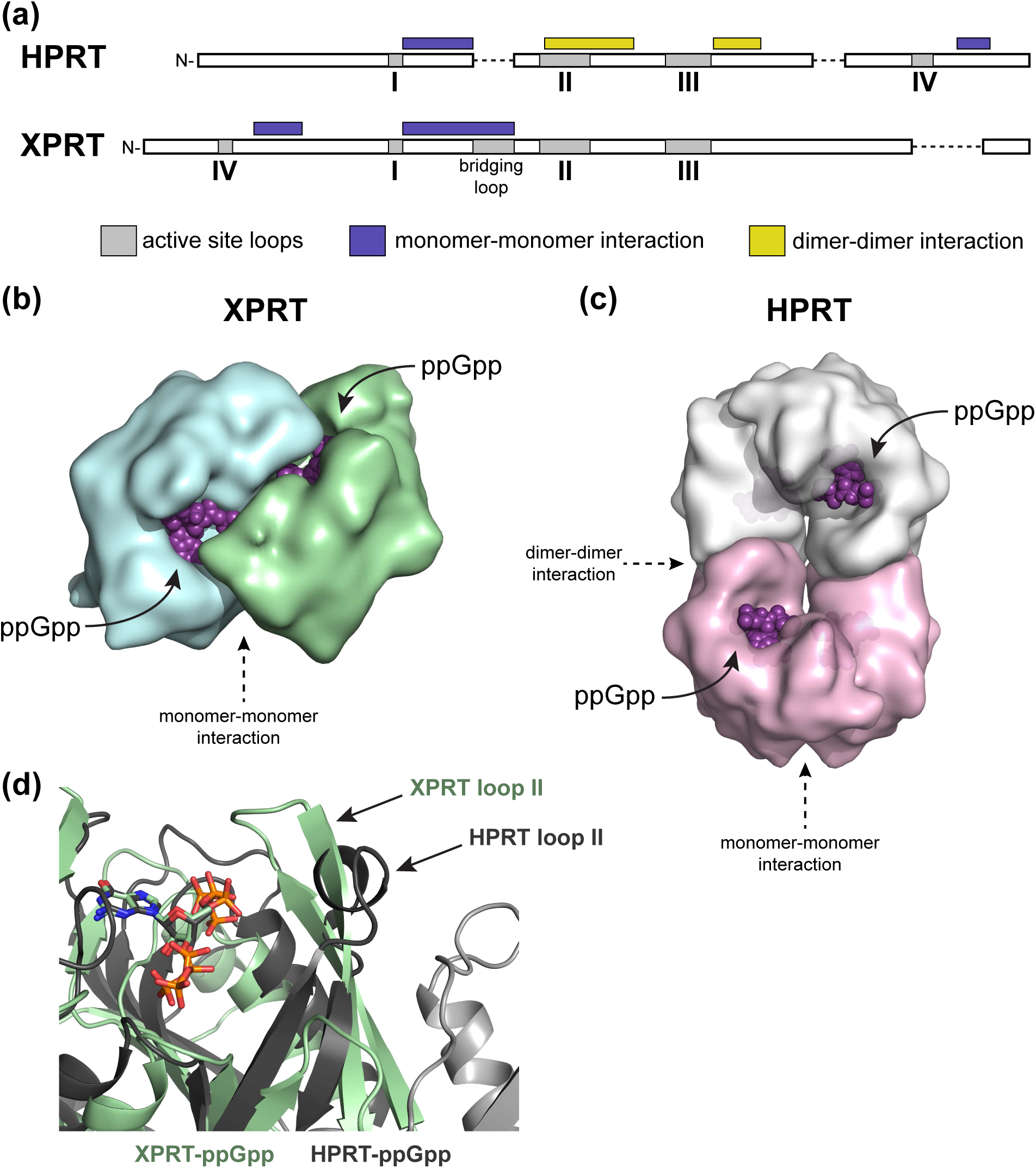
XPRT is allosterically regulated by ppGpp differently than HPRT due to distinct oligomeric interactions. **(a)** Schematic of HPRT and XPRT architecture. Protein structures aligned with PROMALS3D. Gray boxes represent active site loops (I-IV). Regions marked with purple and yellow boxes are involved in monomer-monomer and dimer-dimer interactions, respectively. Dashed lines indicate a break in alignment. **(b)** Surface representation of an XPRT dimer bound to ppGpp. Note the intersubunit binding site that allows XPRT to cooperatively bind ppGpp. **(c)** Surface representation of an HPRT tetramer bound to ppGpp (PDB ID 6D9S). The dimer-dimer interaction allosterically affects the conformation of the ppGpp binding pocket. **(d)** Overlay of XPRT-ppGpp (green) and HPRT-ppGpp (gray) showing the difference in the positioning of loop II. Loop II in HPRT forms a dimer-dimer interaction that holds it away from the binding pocket. Light gray cartoon represents an additional HPRT subunit.

In one case, XPRT displays cooperative binding with (p)ppGpp likely due to a loop at its monomer-monomer interface. The monomer-monomer interfaces of HPRT and XPRT share similar overall secondary structure in an α-helix-turn-β-strand between loops I and II (Figure 8A). However, the sequence identity at this interface has diversified, and, moreover, XPRT has an extra 10-residue loop not found in HPRT (Figure 8A and S2). The (p)ppGpp binding sites in XPRT face one another across the monomer-monomer interface (Figure 8B), and each monomer contributes a loop that bridges the interface to interact with (p)ppGpp and form an electrostatic network of interactions. The bridging loop connects the monomers through (p)ppGpp, providing a likely mechanism for cooperativity we observed between XPRT and (p)ppGpp. In HPRT, on the other hand, the (p)ppGpp binding sites are rotated away from one another on opposite sides of an HPRT dimer (Figure 8C). HPRT lacks the bridging loop and cooperativity binding of ppGpp was not observed ^6^. The electrostatic interactions provided by Arg80 in the bridging loop in XPRT are instead provided by Arg165 in HPRT from within same monomer ^6^. The bridging loop is likely critical for XPRT catalysis as well as inhibition, as it was previously suggested that Arg80 would interact with the substrate PRPP ^11,21^. Although structural information of XPRT bound to PRPP is not available, its interaction with ppGpp in our structure suggests that this would be the case for PRPP as well.

Specificity and affinity of (p)ppGpp binding are also affected by diversification of interfaces between HPRT and XPRT. HPRT has an additional dimer-dimer interface that is responsible for exceedingly tight regulation of HPRT by (p)ppGpp, since the dimer interface sequesters loop II away from the binding pocket to favor inhibitor over substrate binding (Figure 8A and 8D) ^6^. In XPRT, loop II is not held by a dimer-dimer interface. Instead, it partially covers the active site, where it structurally favors alarmones with fewer 5′-phosphates, making pGpp and ppGpp more potent inhibitors of XPRT than pppGpp (Figure 4E). The weakened binding with pppGpp may be due to the energetics required to push loop II away from this site to fit the extra phosphate. The physiological consequences of this loop II conformation are unclear and may be due to selection for enhanced binding of PRPP with its 5′-monophosphate or due to a difference in *in vivo* roles of ppGpp and pppGpp as suggested in *E. coli* ^22^. Regardless, it is clear that the difference in oligomeric interfaces affects affinity and specificity of inhibitor binding.

Overall, elucidating the molecular mechanism of regulation of XPRT has highlighted the strong impact of oligomeric interaction on the evolution of diversification of enzyme regulation, affecting cooperativity, affinity, and specificity of ligand binding.

## MATERIALS AND METHODS

### Determination of XPRT-ppGpp costructure

The coordinate and structure factor files for PDB ID 1Y0B were downloaded from the PDB website. Electron density maps (2F_o_-F_c_ and F_o_-F_c_) were calculated and displayed in COOT ^23^; these clearly showed density for the 5′ portion of ppGpp that was not fit in the deposited structure. Real space refinement in COOT gave acceptable fits in each of the 4 instances of ppGpp. Phenix refine ^24^ was used to refine the structure with the full ppGpp to R_free_ = 15.3 and R_work_ = 20.8. The fit of ppGpp was further examined in COOT. The sodium sites in the 3′ phosphates were checked with CheckMyMetal ^25^ and are not magnesium sites. Bond lengths to the water in between the 5′ phosphates indicate the possibility that a sodium ion binds the 5′ phosphates.

### Plasmid construction and mutagenesis

Plasmids for expression of XPRT for purification were constructed by ligation independent cloning into the pLIC-trPC-HA expression vector (Stols et al., 2002). The *xpt* gene was cloned from *B. subtilis* NCIB 3610 (GenBank accession no. **<u>CP020102</u>**). XPRT variants were created using the QuikChange Site-directed mutagenesis kit according to the manufacturer’s protocol (Agilent). Mutations were confirmed by DNA sequencing. See Table 3 for plasmids, primers, and strains.

**Table 3.**
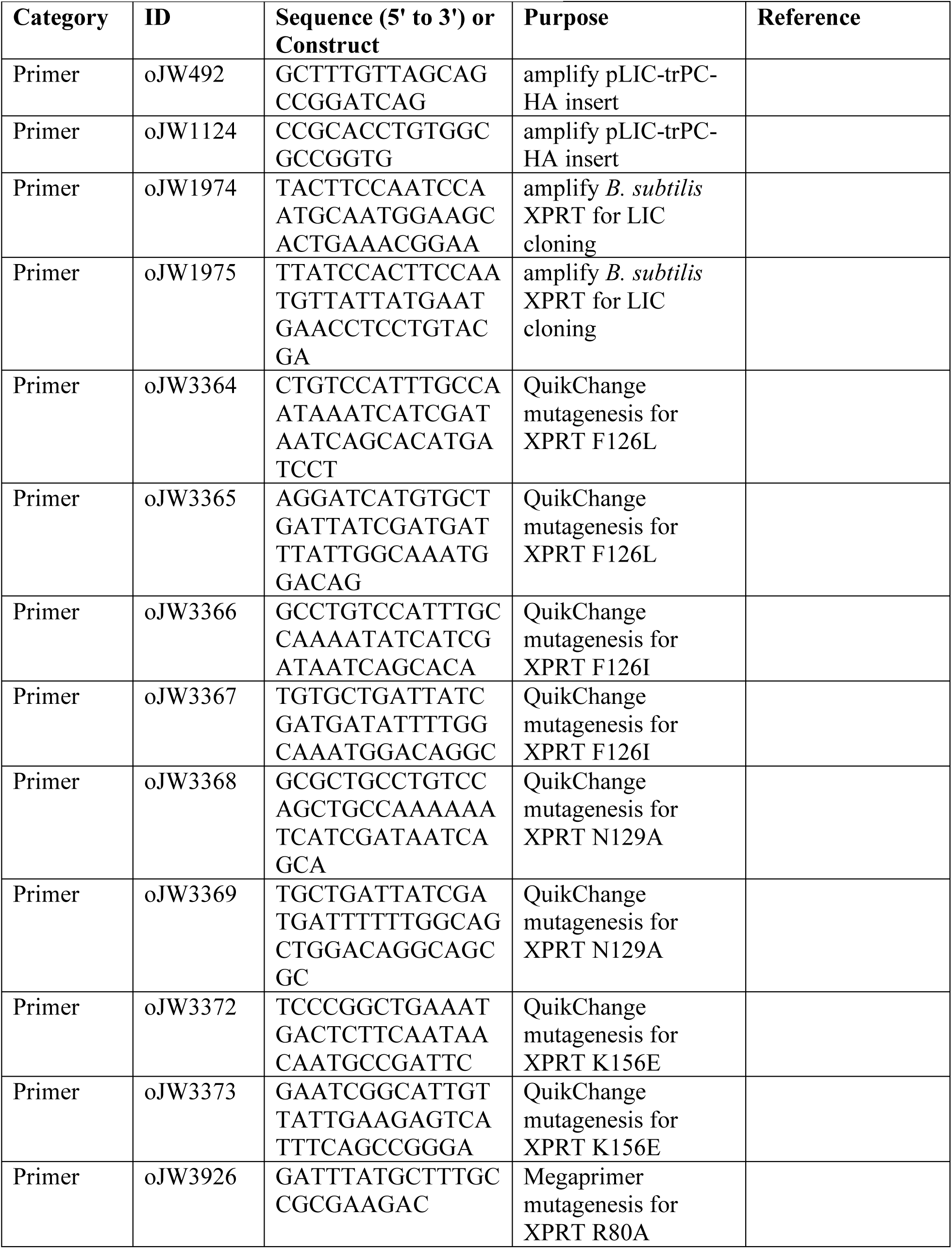
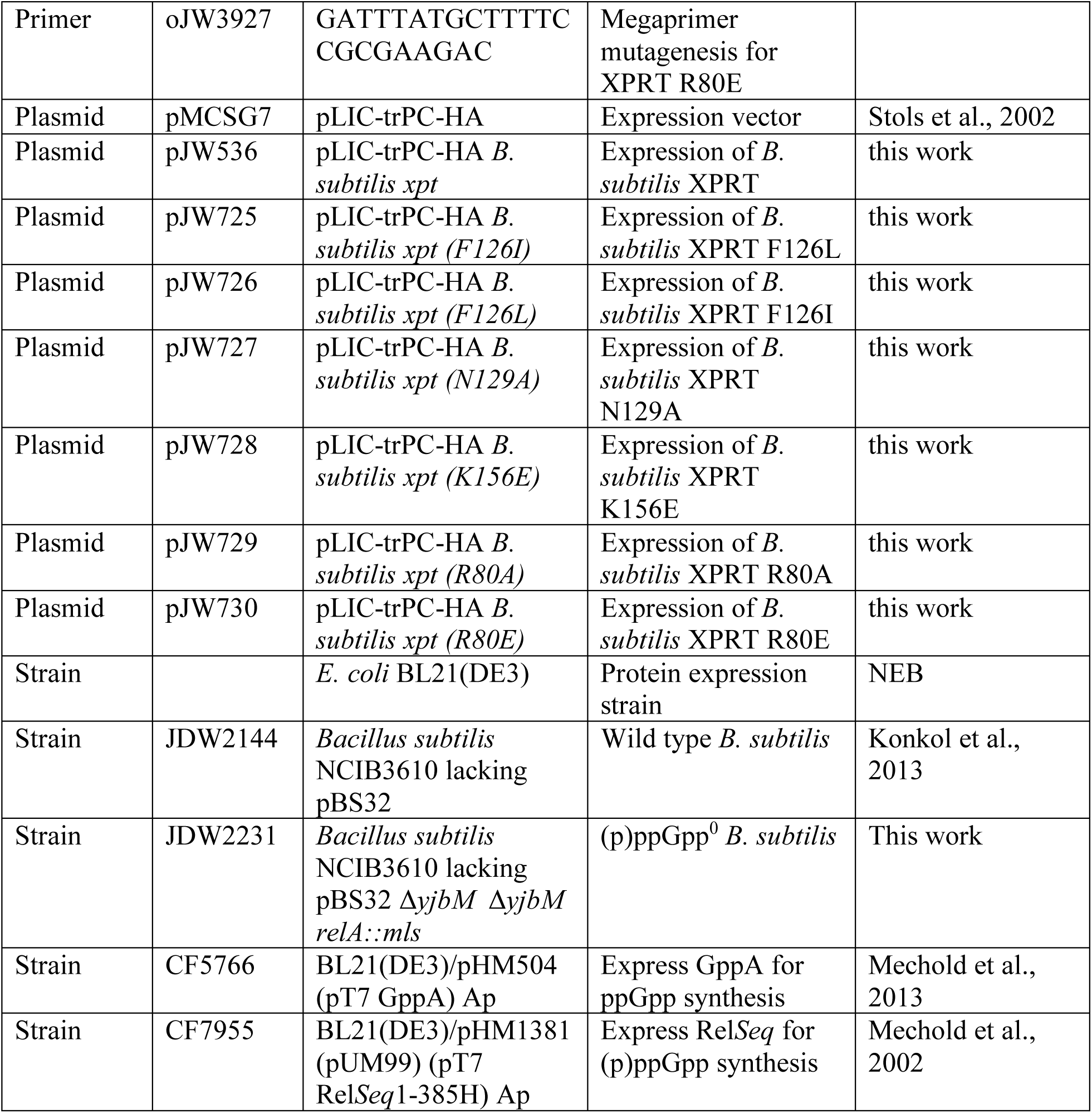
List of primers, plasmids, and strains

### Protein purification

XPRT proteins were purified following a similar protocol ^11^. XPRT was expressed recombinantly in *E. coli* BL21(DE3). A seed culture in LB supplemented with 100 μg/mL carbenicillin (OD ∼ 0.5) was diluted 1:50 into a batch culture of LB medium with carbenicillin. The batch culture was induced with 0.5 mM IPTG at OD_600_ ∼ 0.5 for 3 hours. Cells were pelleted by centrifugation at 10,000 x*g* for 30 min and the pellets were stored at -80 °C. Cell pellets were resuspended in buffer A [50 mM Na_2_HPO_4_, 150 mM NaCl, 20 mM imidazole, pH 8.0] and cells were lysed by French press. Cell debris was removed by centrifuging at 17,000 x*g* for 20 min at 4 °C. The supernatant was filtered, injected into a ÄKTA FPLC (GE Healthcare), and passed over a HisTrap FF column (GE Healthcare). XPRT was eluted with a gradient of the elution buffer [50 mM Na_2_HP0_4_, 150 mM NaCl, 200 mM imidazole, pH 8.0]. Fractions containing purified XPRT were determined via SDS-PAGE analysis. Fractions were collected and dialyzed into 25 mM Tris-HCl (pH 7.6) and 0.1 mM EDTA with two dialysis passages in Spectra/Por dialysis tubing (Spectrum). The concentration of purified XPRT were measured by the Bradford assay (Bio-Rad). The protein was diluted with 20% glycerol, flash frozen, and stored at -80 °C. XPRT lacking the hexahistidine tag was purified for size exclusion chromatography by dialyzing the protein overnight with tobacco etch virus protease in 20 mM Tris-HCl (pH 7.5), 100 mM NaCl, 1 mM DTT, and 0.5 mM EDTA. The protein was dialyzed back to lysis buffer, passaged over a HisTrap FF column, and the flowthrough was collected and concentrated prior to freezing.

XPRT variants were overexpressed as above, but at 10 mL volumes for smaller scale purification. Smaller scale purification was performed with Ni-NTA spin columns (Qiagen). Cell pellets were resuspended in lysis buffer [20 mM sodium phosphate pH 8.0, 500 mM NaCl, 10 mM imidazole] with 1 mg/mL lysozyme and 375 U Benzonase endonuclease (MilliporeSigma), and cells were incubated for one hour on ice for lysis. The lysate was centrifuged at 20,000 x*g* for 15 min. The protein was purified according to the manufacturer’s protocol, with the minor modification of three wash steps and three elution steps. The wash and elution buffers were identical to the lysis buffer, except with 40 mM imidazole and 500 mM imidazole, respectively. Protein purity was determined via SDS-PAGE, protein was dialyzed into 20 mM Tris-HCl pH 7.5, 100 mM NaCl, and 0.1 mM EDTA with Slide-A-Lyzer dialysis devices (ThermoScientific). Protein was concentrated with Amicon Ultra-0.5 concentrators (MilliporeSigma), concentration was determined with the Bradford assay, and protein was frozen with liquid nitrogen for storage at -80 °C.

### XPRT activity assays

XPRT activity assays were performed as previously described ^11^. Briefly, reactions were carried out at 25 °C in a 100 μl mix containing 50 mM Tris-HCl (pH 7.6), 10 mM MgCl_2_, 100 μM xanthine, 1 mM PRPP and 20 nM hexahistidine-tagged XPRT. Reactions were initiated by addition of xanthine. Production of XMP was detected by monitoring absorbance at 252 nm for 10 min in a UV-2401PC spectrophotometer (Shimadzu). A difference in extinction coefficients of 5350 M^-1^cm^-1^ was used for XMP and xanthine to convert absorbance to moles of product. For inhibition curves, assays were performed at the substrate concentrations listed above and at variable pppGpp, ppGpp, and pGpp concentrations. (p)ppGpp was synthesized as described ^26^ and pGpp was a gift from José Lemos. Initial velocities of the inhibited reactions were normalized to the uninhibited initial velocity prior to fitting to the equation Y = 1/(1 + (x / IC_50_)^s^) to calculate IC_50_. For inhibition assays, 50 μM xanthine and 1 mM PRPP were used with varied pppGpp concentrations. For kinetic assays, 100 μM xanthine was used with varied pppGpp and PRPP concentrations. Data fitting was performed using GraphPad Prism.

### Radiolabeled ^32^P-(p)ppGpp synthesis and purification

[5′-α-^32^P]-pppGpp was synthesized from [α-^32^P]-GTP (Perkin Elmer) and ATP using Rel_*Seq*_ (1-385) as previously described ^6,27^. It was purified using a HiTrap QFF anion exchange column (GE Healthcare) as described ^6^. To synthesize [5′-α-^32^P]-ppGpp, 75 mM NH_4_Cl from a 4 M stock was added to a completed ^32^P-pppGpp reaction followed by the addition of 37.5 μg/mL *E. coli* GppA (GppA purified according to ^22^). The reaction was continued for an additional hour and was then treated like a ^32^P-pppGpp reaction for anion exchange purification.

### DRaCALA

Differential radial capillary action of ligand assay (DRaCALA) utilizes the ability of nitrocellulose to separate free ligand from the protein-ligand complex and was used to detect and quantify direct binding of (p)ppGpp to XPRT ^8^. Binding assays were performed on purified hexahistidine-tagged *B. subtilis* XPRT and XPRT variants similarly as described before ^6^. Reactions were carried out in a 15-20 μL reaction mixture with the specified concentration of protein diluted in 20 mM HEPES pH 8, 100 mM NaCl, and 10 mM MgCl_2_. Radiolabeled (p)ppGpp was added at a final concentration of 1:50-1:100 of the first elution fraction from synthesis. Reactions were pipetted or shaken to mix, incubated at room temperature for 10 min, and 2 μL of each reaction was spotted onto Protran BA85 nitrocellulose (GE Healthcare) via pipette or a replicator pinning tool (V&P Scientific, Inc.). After drying for at least 20 min, the nitrocellulose was exposed to a phosphorscreen and scanned with a Typhoon phosphorimager (GE Healthcare, Inc.). Spot intensity was quantified using ImageJ software. Fraction bound ^32^P- (p)ppGpp was calculated and edge effect was corrected as previously described ^8^. Data were analyzed in GraphPad Prism v5.02 and binding curves were fitted to the equation Y = (B_max_ × X^h^) / (K_d_^h^ + X^h^), where h is the Hill coefficient.

### *B. subtilis* viability assays

The *B. subtilis* strains used in this study are NCIB 3610 strain background lacking the pBS32 megaplasmid. Wild type (JDW2144) and (p)ppGpp^0^ *B. subtilis* (JDW2231) were grown on lysogeny broth (LB) and modified Spizizen minimal agar plates (1X Spizizen salts ^28^, 1% glucose, 2 mM MgCl_2_, 0.7 mM CaCl_2_, 50 mM MnCl_2_, 5 mM FeCl_3_, 1 mM ZnCl_2_, 2 mM thiamine, 1.5% agar, and 0.1% glutamate). Since (p)ppGpp^0^ cannot grow on minimal medium, the six amino acids valine, isoleucine, leucine, methionine, threonine, and histidine were included where necessary at concentrations previously reported ^29^. Liquid cultures of *B. subtilis* were grown in LB or S7 defined minimal medium with 1% glucose ^29,30^.

For viability assays, a single colony of each strain was resuspended in S7 medium and spread on modified Spizizen minimal agar plates + 6aa (VILMTH). The strains were grown until small colonies formed (≈16 hrs at 30° C). Cultures were collected from the overnight plates with S7+6aa, diluted to OD_600_ 0.005 and grown in S7+6aa until OD_600_ ≈ 0.2. A pretreatment sample was taken, and the remaining culture was treated with 1 mM xanthine, 1 mM guanine, or water for one hour. Ten-fold dilutions of cultures were plated on LB for quantification of CFU/mL.

### Thin layer chromatography

Thin layer chromatography was performed as described ^31,32^. Wild type and (p)ppGpp^0^ *B. subtilis* were washed from overnight plates and diluted in limited phosphate (1/10 phosphate concentration) S7 minimal media supplemented with 9 amino acids (VILMTHRWE) at previously reported concentrations ^29^. Cultures were grown until OD_600_ ∼ 0.02 and labeled with 50 μCi/ml ^32^P-orthophosphate (900 mCi/mmol; PerkinElmer). When OD_600_ reached 0.2-0.4, cultures were treated with 1 mM xanthine for 30 min. Nucleotides were extracted by mixing 75 μL of culture with 15 μL of cold 2 N formic acid and incubating on ice for at least 20 min. Extracts were centrifuged at 14,000 x*g* for at least 15 min at 4 °C. Two microliters of supernatant were spotted on PEI cellulose TLC plates (MilliporeSigma) and developed in 0.85 M KH_2_PO_4_ pH 3.4. The TLC plates were then exposed to a phosphorscreen and scanned by a Typhoon phosphorimager.

### Alignment

XPRT proteins were selected from representative bacterial species by searching for EC 2.4.2.22 in UniProt. Proteins were chosen that were at least 190 residues since the *B. subtilis* XPRT homologs are longer than either HPRTs (∼180 aa) or XGPRTs (∼150 aa). Proteins were aligned in MEGA X with MUSCLE. (p)ppGpp-interacting residues were determined with LigPlot, and the positions for these residues were selected from the alignment. The frequency logo was generated from the aligned binding residues in WebLogo (https://weblogo.berkeley.edu/logo.cgi). Percent identities between PRT homologs was determined by aligning the proteins in PROMALS3D with PDB ID **<u>1Y0B</u>**, PDB ID **<u>6D9S</u>**, and PDB ID **<u>1A97</u>** used for *B. subtilis* XPRT, *B. subtilis* HPRT, and *E. coli* XGPRT, respectively ^33^. Percent identities were calculated as fraction of identical residues across the whole alignment, including gaps.

### XPRT conservation

16S rRNA sequences for 41 bacterial species representing six phyla were downloaded from NCBI or the Ribosomal Database Project ^34^. 18S rRNA sequences from three eukaryotic species were downloaded from NCBI. Sequences were aligned with CLUSTAL in MEGA X ^35^. MEGA X’s model testing tool was used to select Tamura-Nei model with gamma distributed categories as the best fitting model. A phylogenetic tree was constructed in MEGA X with this model and 100 bootstrap replicates. The presence of XPRT in each species was determined by BLASTing *B. subtilis* XPRT against each species’ proteome. The tree was modified and annotated using the R package ggtree ^36^.

### Differential scanning fluorimetry

Differential scanning fluorimetry was performed as previously described ^37^. Protein (10 μM) was combined with 5X Sypro Orange dye (ThermoFisher, 5000X stock) in a buffer diluted from a 5X stock (final 10 mM HEPES pH 8, 100 mM NaCl, and 10 mM MgCl_2_). The protein and dye were incubated at 25 °C to 90 °C (1 °C/min ramp) using a Bio-Rad CFX Connect Real-time thermocycler. Measurements were taken every minute. T_m_ values were determined as the minimum of the first derivative of the melting curves as provided by the CFX Manager software.

### Size exclusion chromatography

Size exclusion chromatography was performed with an AktaPure FPLC and Superose 12 10/300 GL column (GE Healthcare) with a flow rate of 0.25 mL/min. The mobile phase was 20 mM HEPES pH 8, 300 mM NaCl, and 10 mM MgCl_2_. ppGpp was added to the mobile phase at a final concentration of 35 μM when needed. The column was equilibrated with at least one column volume of buffer prior to addition of protein. The predicted dimer and monomer retention volumes were calculated from the molecular weight of untagged XPRT using a previously published standard curve for this column ^6^ [log(molecular weight) = -0.2575 × retention volume + 7.821]. With a molecular weight of 21038 Da, the predicted retention volumes are 13.58 for an XPRT monomer and 12.41 for an XPRT dimer. The A_280_ (mAU) curves were normalized to have a baseline near zero.

## ACCESSION NUMBERS

Datasets used in this study: PDB ID: 1Y0B, PDB ID: 6D9S, PDB ID: 6D9R, GenBank accession no. CP020102 for *B. subtilis* NCIB 3610 genome. Coordinates and structure factors have been deposited in the Protein Data Bank with accession number 6W1I.

## ACKNOWLEDGMENTS

PDB ID **<u>1Y0B</u>** was deposited by the Midwest Center for Structural Genomics, which was funded by NIH GM62414. This work was funded by NIH R35 GM127088 and R01 GM084003 to J. D. W., and B. W. A. was supported by NSF GRFP DGE-1256259.

## FIGURE LEGENDS

**Figure S1.**
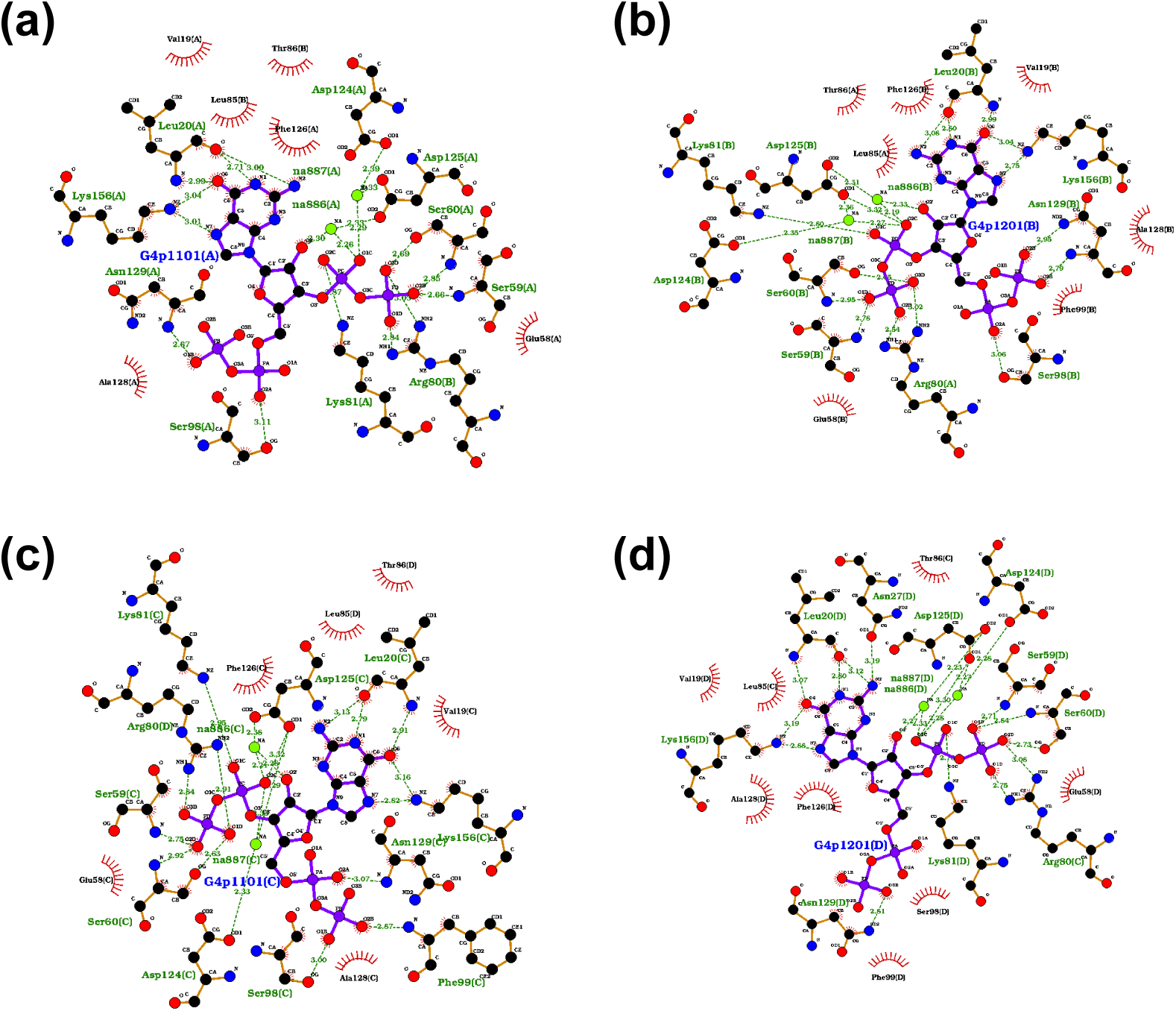
XPRT-ppGpp interactions in the XPRT-ppGpp costructure. Interaction maps of *B. subtilis* XPRT-ppGpp interactions generated by LigPlot+. Each plot represents XPRT-ppGpp interactions for each of the four ppGpp molecules found in the XPRT-ppGpp asymmetric unit. **(a-b)** Interaction maps for two ppGpp molecules in one biological dimer. **(c-d)** Interaction maps for two ppGpp molecules in the second biological dimer.

**Figure S2.**
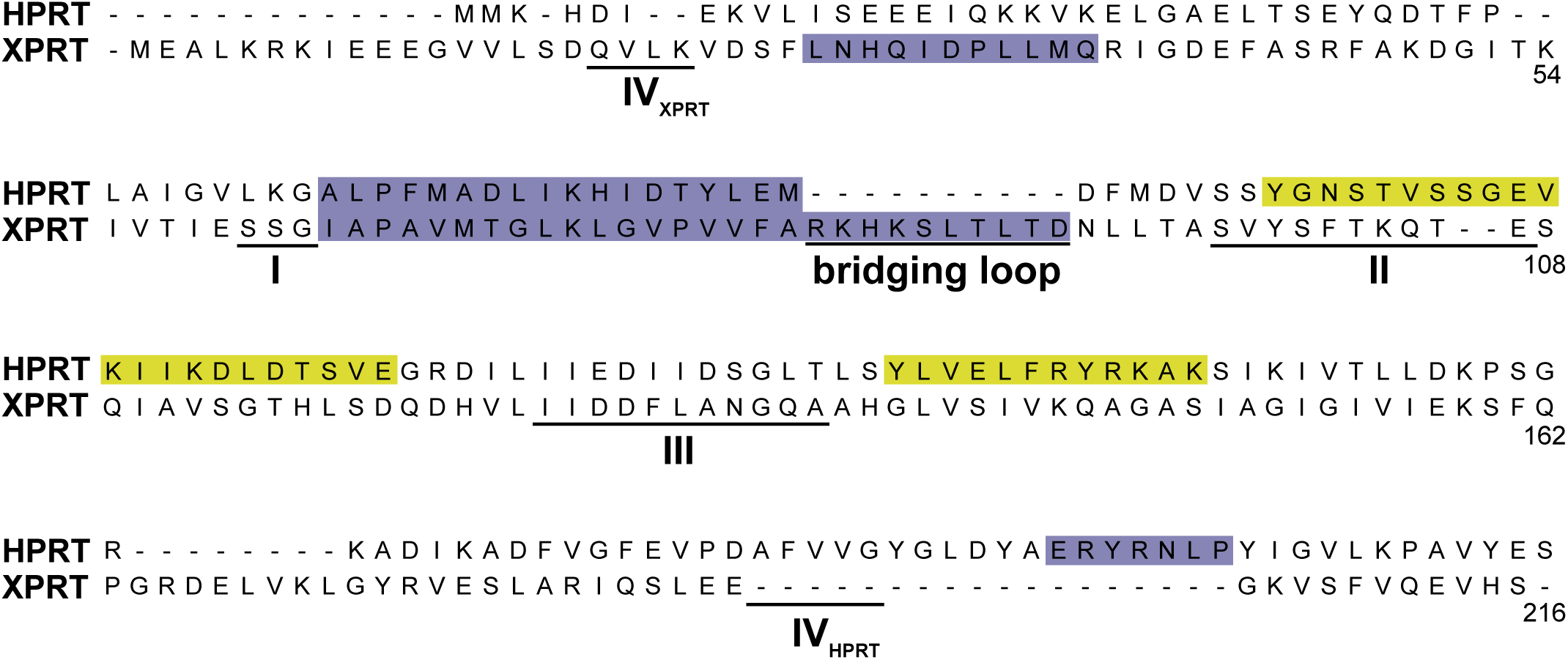
Primary sequence alignment of *B. subtilis* HPRT and XPRT. *B. subtilis* HPRT and XPRT aligned with PROMALS3D with minor manual modifications to the alignment. *B. anthracis* Hpt-1 (PDB ID <u>**6D9S**</u>) and *B. subtilis* XPRT (PDB ID <u>**1Y0B**</u>) were used as the structural models for PROMALS3D. Loops involved in the active site are underlined and numbered I-IV. Regions involved in monomer-monomer interactions are shaded purple. Regions involved in dimer-dimer interactions in HPRT are shaded yellow. Not all shaded residues are directly involved in intersubunit interactions.

## REFERENCES

1. Dakora, F. D. & Phillips, D. a. Root exudates as mediators of mineral acquisition in low-nutrient environments. Plant Soil 245, 35–47 (2002).

2. Traut, T. W. Physiological concentrations of purines and pyrimidines. Mol. Cell. Biochem. 140, 1–22 (1994).

3. Rolfes, R. J. Regulation of purine nucleotide biosynthesis: in yeast and beyond. Biochem. Soc. Trans. 34, 786–790 (2006).

4. Nyhan, W. L. Nucleotide synthesis via salvage pathway. in eLS 1–7 (John Wiley & Sons, Inc., 2014). doi:10.1038/npg.els.0003909

5. Kriel, A. et al. Direct regulation of GTP homeostasis by (p)ppGpp: a critical component of viability and stress resistance. Mol. Cell 48, 231–41 (2012).

6. Anderson, B. W. et al. Evolution of (p)ppGpp-HPRT regulation through diversification of an allosteric oligomeric interaction. Elife 8, e47534 (2019).

7. Christiansen, L. C., Schou, S., Nygaard, P. & Saxild, H. H. Xanthine metabolism in *Bacillus subtilis*: Characterization of the *xpt-pbuX* operon and evidence for purine- and nitrogen-controlled expression of genes involved in xanthine salvage and catabolism. J. Bacteriol. 179, 2540–2550 (1997).

8. Roelofs, K. G., Wang, J., Sintim, H. O. & Lee, V. T. Differential radial capillary action of ligand assay for high-throughput detection of protein-metabolite interactions. Proc. Natl. Acad. Sci. U. S. A. 108, 15528–33 (2011).

9. Gaca, A. O. et al. From (p)ppGpp to (pp)pGpp: characterization of regulatory effects of pGpp synthesized by the small alarmone synthetase of *Enterococcus faecalis*. J. Bacteriol. JB.00324-15 (2015). doi:10.1128/JB.00324-15

10. Krissinel, E. & Henrick, K. Inference of macromolecular assemblies from crystalline state. J. Mol. Biol. 372, 774–797 (2007).

11. Arent, S., Kadziola, A., Larsen, S., Neuhard, J. & Jensen, K. F. The extraordinary specificity of xanthine phosphoribosyltransferase from *Bacillus subtilis* elucidated by reaction kinetics, ligand binding, and crystallography. Biochemistry 45, 6615–27 (2006).

12. Maass, S. et al. Efficient, global-scale quantification of absolute protein amounts by integration of targeted mass spectrometry and two-dimensional gel-based proteomics. Anal. Chem. 83, 2677–2684 (2011).

13. Maass, S. et al. Highly precise quantification of protein molecules per cell during stress and starvation responses in *Bacillus subtilis*. Mol. Cell. Proteomics 13, 2260–2276 (2014).

14. Muntel, J. et al. Comprehensive absolute quantification of the cytosolic proteome of *Bacillus subtilis* by data independent, parallel fragmentation in liquid chromatography/mass spectrometry (LC/MSE). Mol. Cell. Proteomics 13, 1008–1019 (2014).

15. Rymer, R. U. et al. Binding mechanism of metal-NTP substrates and stringent-response alarmones to bacterial DnaG-type primases. Structure 20, 1478–1489 (2012).

16. Bennison, D. J., Irving, S. E. & Corrigan, R. M. The Impact of the Stringent Response on TRAFAC GTPases and Prokaryotic Ribosome Assembly. Cells 8, (2019).

17. Vos, S., Parry, R. J., Burns, M. R., de Jersey, J. & Martin, J. L. Structures of free and complexed forms of *Escherichia coli* xanthine-guanine phosphoribosyltransferase. J. Mol. Biol. 282, 875–889 (1998).

18. Corrigan, R. M., Bellows, L. E., Wood, A. & Gründling, A. ppGpp negatively impacts ribosome assembly affecting growth and antimicrobial tolerance in Gram-positive bacteria. Proc. Natl. Acad. Sci. 113, 201522179 (2016).

19. Zhang, Y., Zbornikova, E., Rejman, D. & Gerdes, K. Novel (p)ppGpp binding and metabolizing proteins of *Escherichia coli*. MBio 9, e02188–17 (2018).

20. Wang, B. et al. Affinity-based capture and identification of protein effectors of the growth regulator ppGpp. Nat. Chem. Biol. 15, 141–150 (2018).

21. Shi, W. et al. Closed site complexes of adenine phosphoribosyltransferase from *Giardia lamblia* reveal a mechanism of ribosyl migration. J. Biol. Chem. 277, 39981–39988 (2002).

22. Mechold, U., Potrykus, K., Murphy, H., Murakami, K. S. & Cashel, M. Differential regulation by ppGpp versus pppGpp in *Escherichia coli*. Nucleic Acids Res. 41, 6175–6189 (2013).

23. Emsley, P. & Cowtan, K. Coot: model-building tools for molecular graphics. Acta Crystallogr. Sect. D Biol. Crystallogr. 60, 2126–2132 (2004).

24. Adams, P. D. et al. PHENIX: a comprehensive Python-based system for macromolecular structure solution. Acta Crystallogr. Sect. D Biol. Crystallogr. 66, 213–221 (2010).

25. Zheng, H. et al. CheckMyMetal: A macromolecular metal-binding validation tool. Acta Crystallogr. Sect. D Struct. Biol. 73, 223–233 (2017).

26. Liu, K. et al. Molecular mechanism and evolution of guanylate kinase regulation by (p)ppGpp. Mol. Cell 57, 735–749 (2015).

27. Mechold, U., Murphy, H., Brown, L. & Cashel, M. Intramolecular regulation of the opposing (p)ppGpp catalytic activities of Rel*Seq*, the Rel/Spo enzyme from *Streptococcus equisimilis*. J. Bacteriol. 184, 2878–2888 (2002).

28. Spizizen, J. Transformation of biochemically deficient strains of *Bacillus subtilis* by deoxyribonucleate. Proc. Natl. Acad. Sci. 44, 1072–1078 (1958).

29. Kriel, A. et al. GTP dysregulation in *Bacillus subtilis* cells lacking (p)ppGpp results in phenotypic amino acid auxotrophy and failure to adapt to nutrient downshift and regulate biosynthesis genes. J. Bacteriol. 196, 189–201 (2014).

30. Vasantha, N. & Freese, E. Enzyme changes during *Bacillus subtilis* sporulation caused by deprivation of guanine nucleotides. J. Bacteriol. 144, 1119–1125 (1980).

31. Bittner, A. N., Kriel, A. & Wang, J. D. Lowering GTP level increases survival of amino acid starvation but slows growth rate for *Bacillus subtilis* cells lacking (p)ppGpp. J. Bacteriol. 196, 2067–76 (2014).

32. Schneider, D. A., Murray, H. D. & Gourse, R. L. Measuring control of transcription initiation by changing concentrations of nucleotides and their derivatives. Methods Enzymol. 370, 606–617 (2003).

33. Pei, J., Kim, B. & Grishin, N. V. PROMALS3D: a tool for multiple protein sequence and structure alignments. Nucleic Acids Res. 36, 2295–2300 (2008).

34. Cole, J. R. et al. Ribosomal Database Project: data and tools for high throughput rRNA analysis. Nucleic Acids Res. 42, 633–642 (2014).

35. Kumar, S., Stecher, G., Li, M., Knyaz, C. & Tamura, K. MEGA X: Molecular evolutionary genetics analysis across computing platforms. Mol. Biol. Evol. 35, 1547–1549 (2018).

36. Yu, G., Smith, D. K., Zhu, H., Guan, Y. & Lam, T. T. Y. GGTREE: an R package for visualization and annotation of phylogenetic trees with their covariates and other associated data. Methods Ecol. Evol. 8, 28–36 (2017).

37. Niesen, F. H., Berglund, H. & Vedadi, M. The use of differential scanning fluorimetry to detect ligand interactions that promote protein stability. Nat. Protoc. 2, 2212–2221 (2007).

